# Analyzing the Involvement of Cholinesterases in the Immune Landscape of Lung Adenocarcinoma and Their Prognostic Values

**DOI:** 10.1101/2024.05.09.593422

**Authors:** Fengyu Zhang, Zhouhai Zhu, Ying Guan, Meng Li, Zhenhua Pan, Ju Wang

**Author notes:** These two authors contributed equally to this work (Fengyu Zhang, Zhouhai Zhu). Corresponding authors. Correspondence to: Ju Wang, PhD.

## Abstract

Lung adenocarcinoma (LUAD) is the most common subtype of lung cancer and its prognosis is poor. The cholinergic system is involved in the development of lung cancer but its role is still unclear. In this study, we collected 231 cholinergic-related genes, and examined their expression in LUAD samples and normal tissues, from which 37 differentially expressed genes were screened. Then, by survival analysis, we identified 7 genes related to the prognosis of LUAD, among which acetylcholinesterase (ACHE) and butyrylcholinesterase (BCHE) were included. The expression of AChE was upregulated in LUAD samples, and its expression had a significant positive correlation with the prognosis of male patients. But the expression of BChE was down-regulated in LUAD samples, and the elevated BChE expression was associated with a good prognosis in women and non-smoking patients. We also observed a close relationship between the two genes and immune landscape of LUAD. The AChE high expression patients had a higher ratio of tumor-infiltrating immune cells than the low expression patients, while the BChE high expression group had higher ratios of both tumor-infiltrating immune cells and stromal cells. We collected a total of 113 immunomodulatory genes associated with AChE and BChE to build an immunoregulatory network, which comprised several gene clusters. We also found that the expression of AChE and BChE was associated with immune escape in LUAD. Our results showed that AChE and BChE may play an important role in the development of LUAD, and could be promising biomarkers and targets for its diagnosis and treatment.

## 1. Introduction

Lung cancer is one of the most common cancers worldwide. With an estimated 2 million new cases and 1.8 million deaths per year, lung cancer claims more lives than breast, colon and prostate cancers combined, and accounts for about one-fifth of all cancer-related deaths (Bade and Dela Cruz 2020). In the past decades, we have made much progress in diagnosing and treating lung cancer due to improvements in our understanding on the biological features of the disease, application of predictive biomarkers and refinements in its treatment (Howlader et al. 2020). Despite better outcomes can be achieved as a result of increasingly sophisticated combinations of surgery and different therapies in general, the survival rate of lung cancer is still limited (Garon et al. 2019). Therefore, it is necessary to develop new therapeutic approaches for this disease.

In recent years, a number of potential new targets for lung cancer have been reported (Alvarez et al. 2017), among which genes involved in the cholinergic system are included (Profita et al. 2008; Song et al. 2008; Song et al. 2003). The cholinergic system transmits information through acetylcholine (Ach), which is not only a growth factor for lung cancer but also can stimulate the adhesion, migration and invasion of lung cancer cells (Wang et al. 2001). The cholinergic system may participate in lung cancer growth and progression by different mechanisms. One way is the autocrine cholinergic loop. Some cells in lung or related organs, such as bronchial epithelial cells, pulmonary neuroendocrine cells and airway cells, can express nicotinic acetylcholine receptors (nAChRs) and muscarinic acetylcholine receptors (mAChRs); in addition, they can also synthesize and secrete ACh (Minna 2003; Wang et al. 2001). Activation of the receptors by ACh secreted by the tumor cells or by cholinergic agonists like nicotine in tobacco, can stimulate tumor growth (Spindel 2016). Another approach is the tumor-nervous connections, through which the peripheral nerves may modulate the biological behavior of the cancer cells and influence tumor development by releasing neurotransmitters like ACh into the tumor microenvironment and activating the corresponding receptors (Arese et al. 2018; Entschladen et al. 2004). By modulating cholinergic receptors, the cholinergic system can provide functional input to tumors and is involved in the growth, angiogenesis and metastasis of lung cancer (Friedman et al. 2019). Therefore, the cholinergic pathway plays an important role in the occurrence and development of lung cancer; and molecules in this system, including those involved in choline transport, ACh synthesis, secretion, degradation, nicotinic signaling and muscarinic signaling, are promising targets for therapeutic intervention (Spindel 2016).

Smoking is the major cause of lung cancer because nAChRs can be activated by nicotine in cigarettes thereby accelerating the growth of lung cancer. For this reason, of all the components of the cholinergic system, the relationship between nAChRs and lung cancer has been extensively studied. While our understanding on the role of other components, such as cholinesterases, is still incomplete (Grando 2014).

In this study, we investigated the involvement of the cholinergic system in lung adenocarcinoma (LUAD), the most common histological type of lung cancer. More specifically, we focused on cholinesterases, i.e., acetylcholinesterase (AChE) and butyrylcholinesterase (BChE), the proteins responsible for the rapid breakdown of Ach into choline and acetate and causing termination of the excitatory effect of acetylcholine on the postsynaptic membrane. Briefly, we screened the cholinergic genes that are differentially expressed in LUAD tissues, and identified key genes related to prognosis of the disease. We then analyzed the relationship between these genes and immune cells, immune-related pathways, and immunotherapy response in patients with LUAD. We found that cholinesterases can influence the occurrence and development of LUAD by affecting the tumor microenvironment and immune-related pathways, and may be potential therapeutic targets for LUAD therapeutic intervention.

## 2. Materials and methods

### 2.1 Collection of genes involved in the cholinergic system

The genes involved in the cholinergic system were retrieved from PubMed (https://pubmed.ncbi.nlm.nih.gov/) and the major pathway databases, including KEGG (Kanehisa et al. 2021), Reactome (Jassal et al. 2020), Wikipathway (Martens et al. 2021) and Pathway Commons (Rodchenkov et al. 2020). Six cholinergic-related pathways, i.e., cholinergic synapse, choline metabolism in cancer, choline catabolism, acetylcholine neurotransmitter release cycle, choline biosynthesis III, acetylcholine synthesis, were retrieved, from which a total of 231 cholinergic system-related genes were collected.

### 2.2 Identification of genes related to LUAD

The transcriptomic data of LUAD and adjacent normal tissues were downloaded from the Cancer Genome Atlas (TCGA) (https://portal.gdc.cancer.gov/repository). The criteria for inclusion in the TCGA dataset were: (1) primary solid tumor sample of LUAD and solid tissue normal samples; (2) sequencing samples from frozen tissue of patients; and (3) with available survival information. The genome-wide gene expression profiles of 494 LUAD tumor samples and 57 adjacent non-tumor samples measured by RNA-Seq were obtained.

Based on the clinical information of the retrieved samples, patients that smoked or had stopped smoking for less than or equal to 15 years, were classified as smokers, while those did not smoke or had stopped smoking for more than 15 years were classified as nonsmokers (Sui et al. 2020). Because the number of non-smoking patients was relatively small, to reduce the potential bias caused by the unbalanced sample sizes, the transcriptomic dataset GSE31210 in the GEO database was adopted (Okayama et al. 2012). This dataset contained the genome-wide expression profiles of 226 LUAD patients measured by Affymetrix U133Plus2.0 arrays. We retrieved the non-smoking samples from the dataset and merged them with the LUAD tumor samples of TCGA. By this way, 115 samples of non-smoking LUAD patients were collected from the GEO database (GSE31210) to form a new dataset with the LUAD cohort from TCGA. Eventually, a dataset including 57 normal samples and 609 tumor samples was obtained (Supplementary Table S1). To reduce heterogeneity caused by differences between the datasets, we normalized the combined matrix and remove the batch effect using the R package“sva”.

To identify differentially expressed genes (DEGs) between LUAD tissues and adjacent normal tissues, the R package“DESeq2” was used (Love et al. 2014). The DEGs were identified by a false discovery rate (FDR) <0.01, as well absolute logarithmic fold change (| log FC|)> 2.0.

The relationship between DEGs and patients’ prognosis was investigated. Briefly, we utilized Kaplan-Meier curves to compare the overall survival differences between “high” and “low” expression groups of DEGs, and calculated p-value for each gene using the log-rank test in the survival package of R.

The impact of clinical features on patient prognosis was revealed by univariate and multivariate Cox analysis using the R package “survival”. Features of patient gender, age, race, smoking status, and tumor stage were analyzed to identify the independent prognostic features in patients with LUAD (Supplementary Table S2). Next, we divided the sample into two groups by median and used Kaplan-Meier survival curves and log-rank tests to analyze the relationship between key genes expression, gender, smoking status, and prognosis of LUAD patients.

### 2.3 Correlation analysis of key genes and immune infiltration

Then, the correlation between the key genes and immune response was investigated. Briefly, the level of infiltrating stromal cells and immune cells was analyzed by ESTIMATE (https://bioinformatics.mdanderson.org/public-software/estimate/), a tool for predicting tumor purity and the presence of infiltrating stromal/immune cells in tumor tissues by calculating three scores (stromal score, immune score and estimate score) using gene expression data (Yoshihara et al. 2013). It calculates stromal score and immune score by performing single sample gene set enrichment analysis (ssGSEA). The mRNA samples of key genes were divided into high- and low-expression groups according to the median, and the R package “estimate” was used to analyze the relationship between key genes expression differences and tumor purity, and to measure the relationship with immune infiltration.

We further analyzed the correlation between key genes expression and immune cells using TIMER (https://cistrome.shinyapps.io/timer/) and CIBERSORT (https://cibersortx.stanford.edu/). TIMER allows the analysis of key genes in relation to 6 types of immune cells (B cells, CD4+T cells, CD8+T cells, macrophages, neutrophils and dendritic cells) (Li et al. 2017); and CIBERSORT is a deconvolution algorithm to predict the proportion of 22 tumor-infiltrating immune cells in each sample based on gene expression data (Newman et al. 2015).

### 2.4 Building an immunomodulatory network for key genes

Since interactions between tumor cells and immune cells play an important role in both cancer development and treatment, we further built an immunomodulatory network to analyze the roles of the key genes in LUAD. We collected immunoregulatory genes significantly associated with AChE and BChE from the TISIDB database (https://cis.hku.hk/TISIDB/), an online database that integrates data on tumor and immune system interactions (Ru et al. 2019), and then constructed a PPI network of immunoregulatory genes using the STRING database (Szklarczyk et al. 2021). The networks were visualized using Cytoscape software (version 3.9.1) and key modules were explored using the MCODE plugin (Bader and Hogue 2003). Subsequently, the function features of the modules were analyzed by “ClusterProfiler” package in R.

### 2.5 Assessment of response to immune checkpoint blockade (ICB) therapy

The immune checkpoint blockade (ICB) therapy has been applied for cancer treatment and delivered promising clinical outcomes. We analyzed the relationship between AChE and BChE expression and patient response to immunotherapy via the TIDE algorithm (http://tide.dfci.harvard.edu) (Jiang et al. 2018). To predict ICB response, the TIDE algorithm models two mechanisms of tumor immune escape: T cell dysfunction and the prevention of T cell infiltration. The TIDE scores provide a better assessment of the efficacy of anti-PD1 and anti-CTLA4 therapy compared to biomarkers such as PD-L1 levels and tumor mutational load.

## 3. Results

### 3.1 Screening of cholinergic genes involved in LUAD

For the 231 genes involved in the cholinergic system (Supplementary Table S3), we compared their expressions in the LUAD samples and normal tissues, a total of 37 differentially expressed genes were obtained (Supplementary Table S4; Figure S1). Among these genes, there were several subunits of nicotinic or muscarinic acetylcholine receptors (e.g., CHRNA2, CHRNA5, CHRNB4, CHRNA9, CHRND, CHRM1 and CHRM2), a couple genes encoding subunits of G proteins (e.g., GNGT1, GNG4 and GNB3), several member of the potassium channel family (e.g., KCNJ4, KCNJ6, KCNJ18, KCNQ2, KCNQ3 and KCNQ5), membrane-spanning transporters (e.g., SLC18A3 and SLC44A5), and those involved in other functions. Three genes related to metabolism of choline, i.e., CHAT, AChE and BChE, were also included.

The prognostic value of these genes was assessed by survival analysis (Supplementary Figure S2). Of these genes, seven showed significant association with prognosis in LUAD patients. We found that patients with higher expression levels of AChE, BChE, CAMK2B, GNB3, PIP5K1B and PLA2G4F displayed better prognosis, while higher expression of KCNQ3 was associated with worse prognosis.

Among these seven genes, CAMK2B, GNB3, PIP5K1B, PLA2G4F and KCNQ3, are related to various functions, such as calcium signaling, protein phosphorylation, PI3K signaling, glycerophospholipid metabolism, or neuronal excitability. They usually locate at the downstream of the cholinergic system and are not specifically related to the binding or metabolism of acetylcholine. The other two genes, AChE and BChE are hydrolases of acetylcholine and play key roles in the cholinergic system, and were chosen for the subsequent analysis.

### 3.2 Expression patterns of AChE and BChE and the prognosis of LUAD

By analyzing the impact of several factors (i.e., gender, age, race, smoking status, and tumor stage) on prognosis in LUAD patients, gender and smoking status were found to have hazard ratios less than 1.0, indicating they were related to a reduction in the hazard of death in LUAD patients (Supplementary Table S2). We further checked the correlation between the expression levels of AChE and BChE with gender and smoking status in LUAD patients. AChE was upregulated in LUAD samples, and its expression had a significant positive correlation with the prognosis of male patients (P=0.028, Figure 1A). However, there was no significant difference between the expression of AChE and prognosis in female patients, non-smoking patients, and smoking patients (Figure 1B-D). The expression of AChE was higher in both male smoking and male non-smoking patients than in normal tissues (Kruskal-Wallis test, P=1.4×10^-5^), which was also consistent with the up-regulation of AChE expression in LUAD (Figure 1E-F). Since the expression of AChE was higher in male non-smokers than in male smokers, we further analyzed the relationship between AChE and the prognosis of these two types of patients. Actually, the elevated AChE expression was associated with good prognosis, and male non-smoking patients with high AChE expression had the highest survival rate (Figure 1G; P=0.0041).

**Figure 1.**
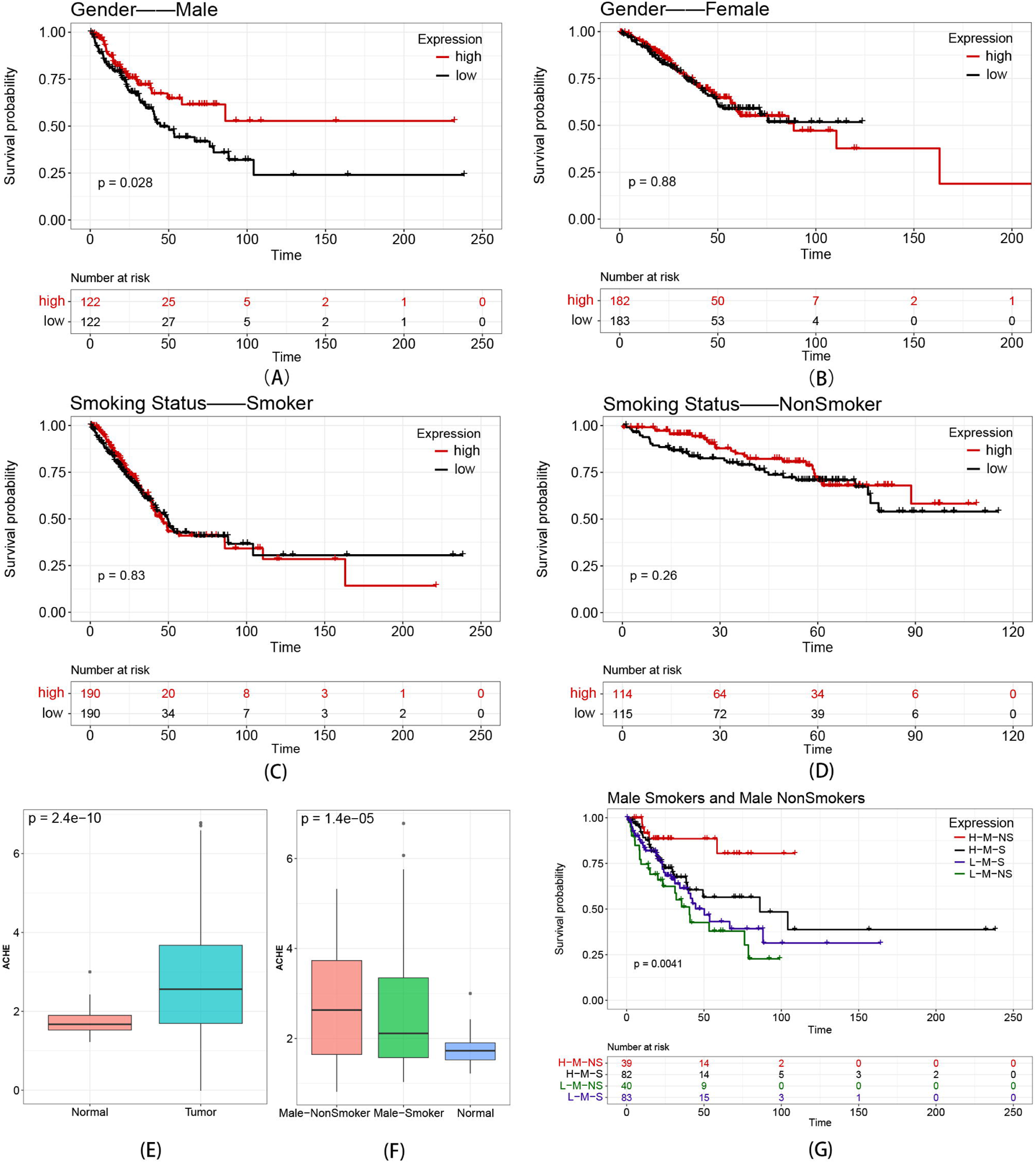
Survival analysis of AChE. (A) Survival curves of AChE in male LUAD patients; (B) survival curves of AChE in female LUAD patients; (C) survival curves of AChE in smoking LUAD patients; (D) survival curves of AChE in non-smoking LUAD patients; (E) the difference in gene expression of AChE in tumor and normal tissues; (F) the differences in gene expression of AChE in male smoking, male non-smoking and normal tissues; (G) relationship between AChE expression and the prognosis of male smoking and male non-smoking LUAD patients.

The expression of BChE was down-regulated in LUAD (Figure 2E). We found that elevated BChE expression was associated with a good prognosis in women and non-smoking patients (Figure 2B and 2D) and was not significantly associated with the prognosis of men, or smoking patients(Figure 2A and 2C). The expression of BChE was lower than that of normal tissues in all four categories of patients (Figure 4F), which is also in accordance with the down-regulated expression of BChE in LUAD. The expression of BChE was higher in female nonsmoking patients than in female smoking patients, male nonsmoking patients, and male smoking patients (Figure 2F). As seen in Figure 2G, female non-smoking patients had a higher survival rate than male non-smoking patients, and female non-smoking patients with high BChE expression presented the best prognosis.

**Figure 2.**
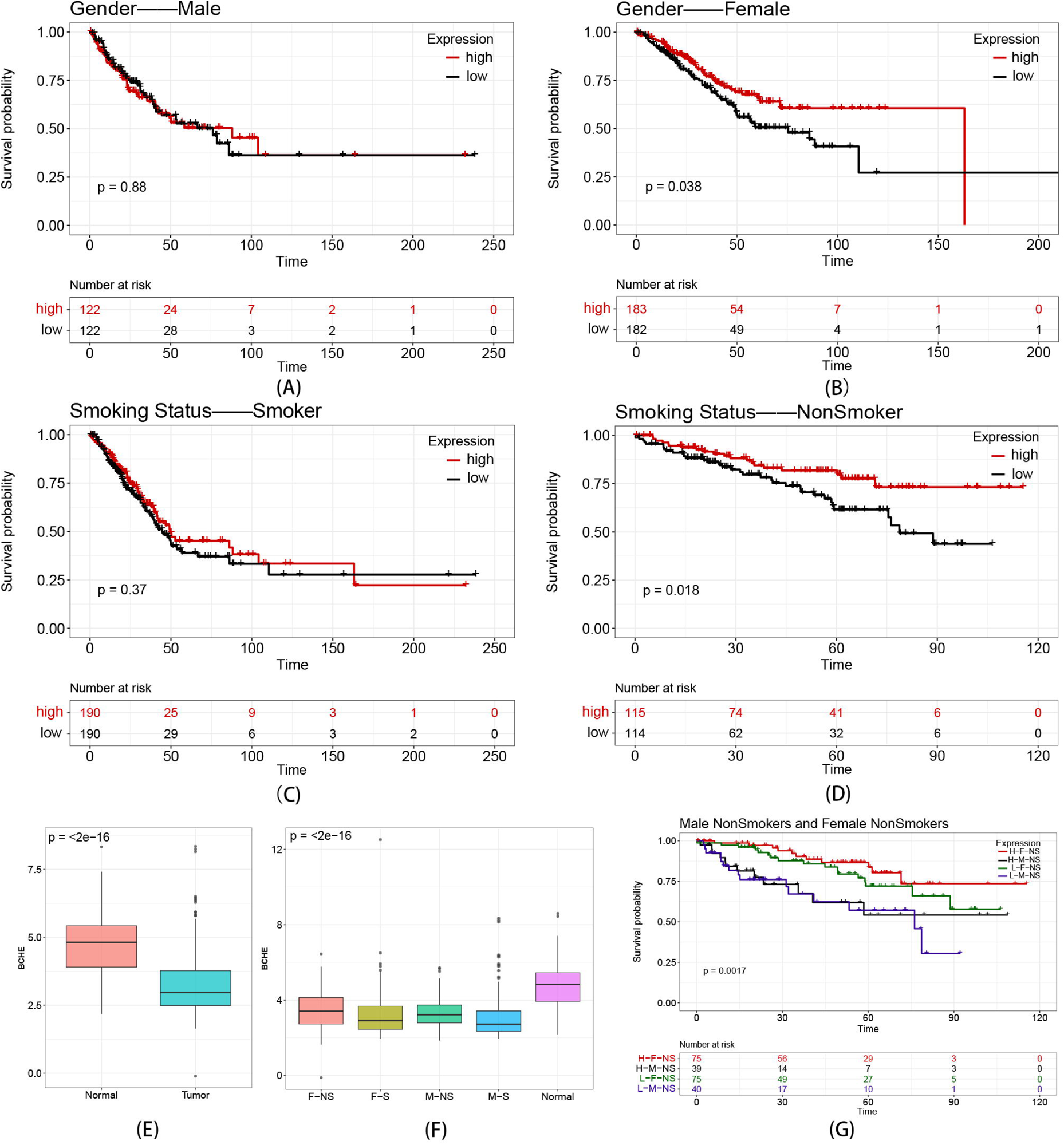
Survival analysis of BChE. (A) Survival curves of BChE in male LUAD patients; (B) survival curves of BChE in female LUAD patients; (C) survival curves of BChE in smoking LUAD patients; (D) survival curves of BChE in non-smoking LUAD patients; (E) the difference in gene expression of BChE in tumor and normal tissues; (F) the differences in BChE expression in female non-smoking, male non-smoking, female smoking, male smoking LUAD patients and normal tissues; (G) relationship between BChE expression and the prognosis of non-smoking LUAD patients.

Thus, AChE and BChE expressed differentially with respect to gender and smoking status. The expression level of AChE was positively correlated with the prognosis of male patients, especially in male non-smoking patients; while elevated expression of BChE was associated with a favorable prognosis in female patients and non-smoking patients, and the best prognosis in female non-smoking patients.

### 3.3 Correlation of AChE and BChE with immune response in LUAD

Our analyses further revealed that both AChE and BChE were related to immune infiltration. By ESTIMATE, we assessed the presence of infiltrating stromal/immune cells in tumor tissues of LUAD using gene expression data of AChE and BChE For AChE, its expression showed a significant difference in the ImmuneScore corresponding to in the high expression group and the low expression group (*p*=0.0049), but not the StromalScore (*p*=0.067) (Supplementary Figure S3A), which indicated that the proportion of tumor-infiltrating immune cells in the two groups was different, with the AChE high expression group had a notably higher ratio of such cells. In addition, the ESTIMATEScore was also different for the two groups (*p*=0.013), indicating the AChE high expression group had lower tumor purity than the low expression group. For BChE, its expression showed a significant difference in the ImmuneScore corresponding to in the high expression group and the low expression group (*p*=0.026), as well as the StromalScore (*p* =2.70×10^-5^) (Supplementary Figure S3B), which indicated that the proportions of tumor-infiltrating immune cells and stromal cells in the two groups were different, with the BChE high expression group had notably higher ratios of both tumor-infiltrating immune cells and stromal cells. In addition, the ESTIMATEScore was also different for the two groups (*p* =9.80×10^-4^), indicating the BChE high expression group had lower tumor purity than the low expression group.

We then analyzed the relationship between the expression level of the two genes and the six types of major immune cells using TIMER. The expression of AChE was positively correlated with B cells (*p* =7.07×10^-8^), CD4+T cells (*p* =2.19×10^-11^), and dendritic cells (*p* =0.0081), and negatively correlated with CD8+T cells (*p* =0.0343) (Supplementary Figure S4A). For BChE, its expression had a significant positive correlation with all the six types of immune cells, i.e., B cells (*p* =0.0189), CD8+T cells (*p* =2.25×10^-4^), CD4+ T cells (*p* =5.61×10^-5^), macrophages (*p* =8.54×10^-15^), neutrophils (*p* =1.71×10^-6^), and dendritic cells (*p* =5.96×10^-7^) (Supplementary Figure S4B).

The relationship between the expression of the two genes and the immune cells was confirmed by the results from the CIBERSORT algorithm, which estimated the abundances of 22 immune cells in the LUAD samples based on the gene expression data. By analyzing the effect of immune cell content on prognosis, we found that the higher proportion of monocytes and M0 macrophages was associated with better prognosis, while higher proportion of M2 macrophages, activated dendritic cells, gamma delta T cells and regulatory T cells was linked with a worse prognosis (Supplementary Figure S5).

Then, the abundances of these 22 immune cells within the high and low expression group of AChE and BChE were compared (Figure 3). The abundance of memory B cells, T follicular helper cells, M0 microphages and M1 microphages, was significantly higher in the AChE high expression group. On the other hand, The immune cells with higher abundances in the AChE low expression group included plasma cells, M2 microphages, activated dendritic cells and Eosinophils. In summary, higher expression of AChE was associated with a higher abundance of M0 microphages, as well as a lower abundance of M2 microphages and activated dendritic cells. At the same time, a higher abundance of M0 microphages and lower abundance of M2 microphages and activated dendritic cells was associated with better prognosis in LUAD patients (Supplementary Figure S5), Since AChE expression was upregulated in tumor tissues of LUAD, it was likely that AChE was associated with the suppression of LUAD. On the other hand, the abundances of resting memory CD4+ T cells, activated NK cells, monocytes, M0 microphages and resting mast cells, were higher in BChE high expression group; and the abundances of regulatory T cells and activated mast cells were higher in the BChE low expression group. Since BChE was downregulated in tumor tissue, it was likely to inhibit tumor growth.

**Figure 3.**
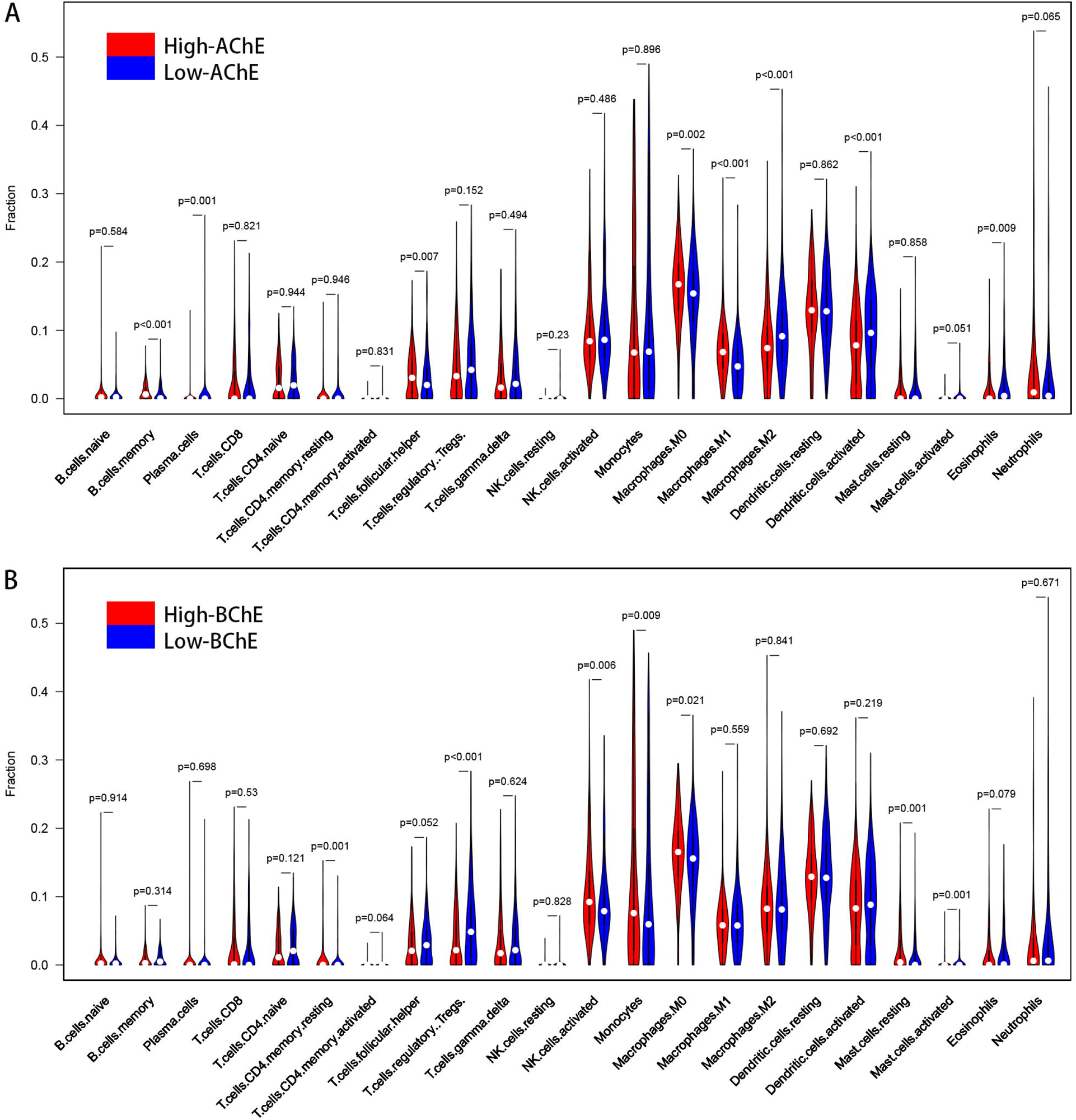
Correlation between the expression levels of AChE and BChE and the level of immune cell infiltration. (A) The difference in the proportion of 22 types of immune cells between the groups with high and low AChE expression. Red and blue symbol represents the high and low AChE expression group, respectively; (B) the difference in the proportion of 22 types of immune cells between the groups with high and low BChE expression. Red and blue symbol represents the high and low AChE expression group, respectively.

### 3.4 Immunoregulatory network of AChE and BChE

We collected a total of 113 immunomodulatory genes associated with AChE and BChE (p<0.05), including chemokines, receptors, MHC, immunostimulators and immunostimulators (Table 1). To investigate the interactions between these genes, we constructed a PPI network composing 113 nodes and 1901 edges (Figure 4). To determine key modules, the network was subdivided into 4 clusters using the MCODE algorithm. Enrichment analysis revealed that the genes in Cluster 1 were mainly involved in pathways like leukocyte chemotaxis, cell chemotaxis, chemokine-mediated signaling, viral protein interaction with cytokine and cytokine receptor, as well as cytokine-cytokine receptor interaction. Genes in Cluster 2 were associated with functions such as regulation of T cell activation, positive regulation of leukocyte cell-cell adhesion, cell adhesion molecules, and cytokine-cytokine receptor interaction. The genes in Cluster 3were closely related to MHC complex and immune response, while those in Cluster 4 genes were involved in the positive regulation of interferon-γ.

**Table 1.**
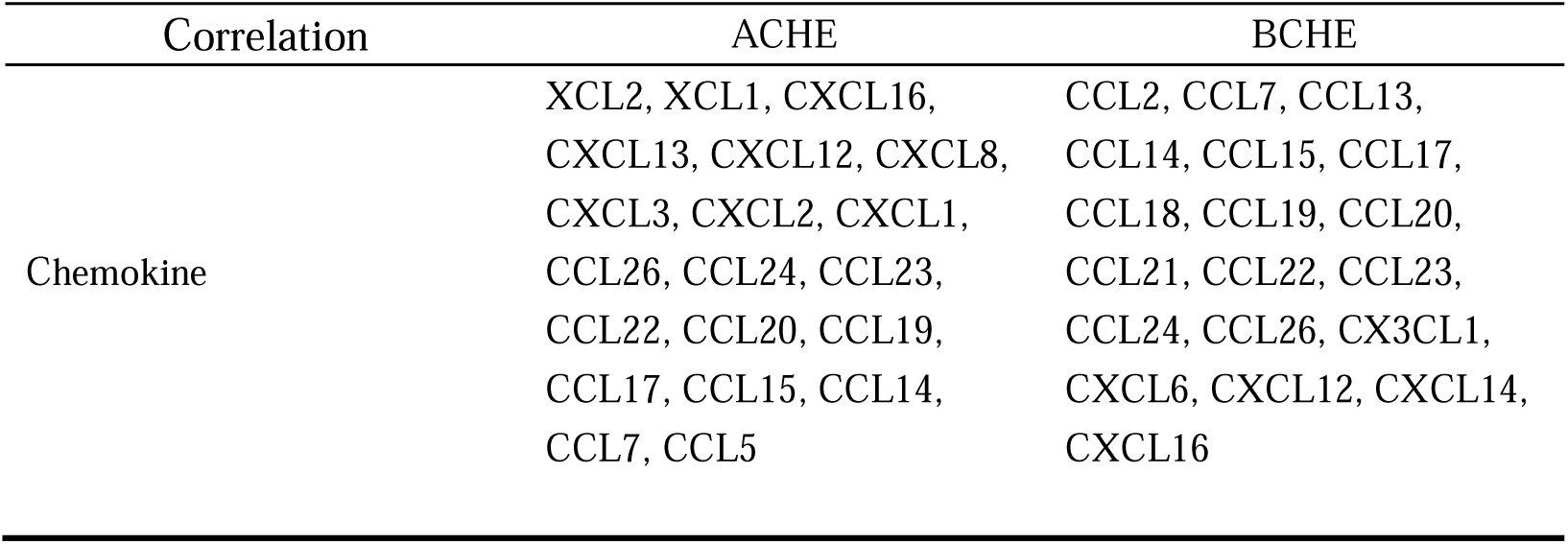
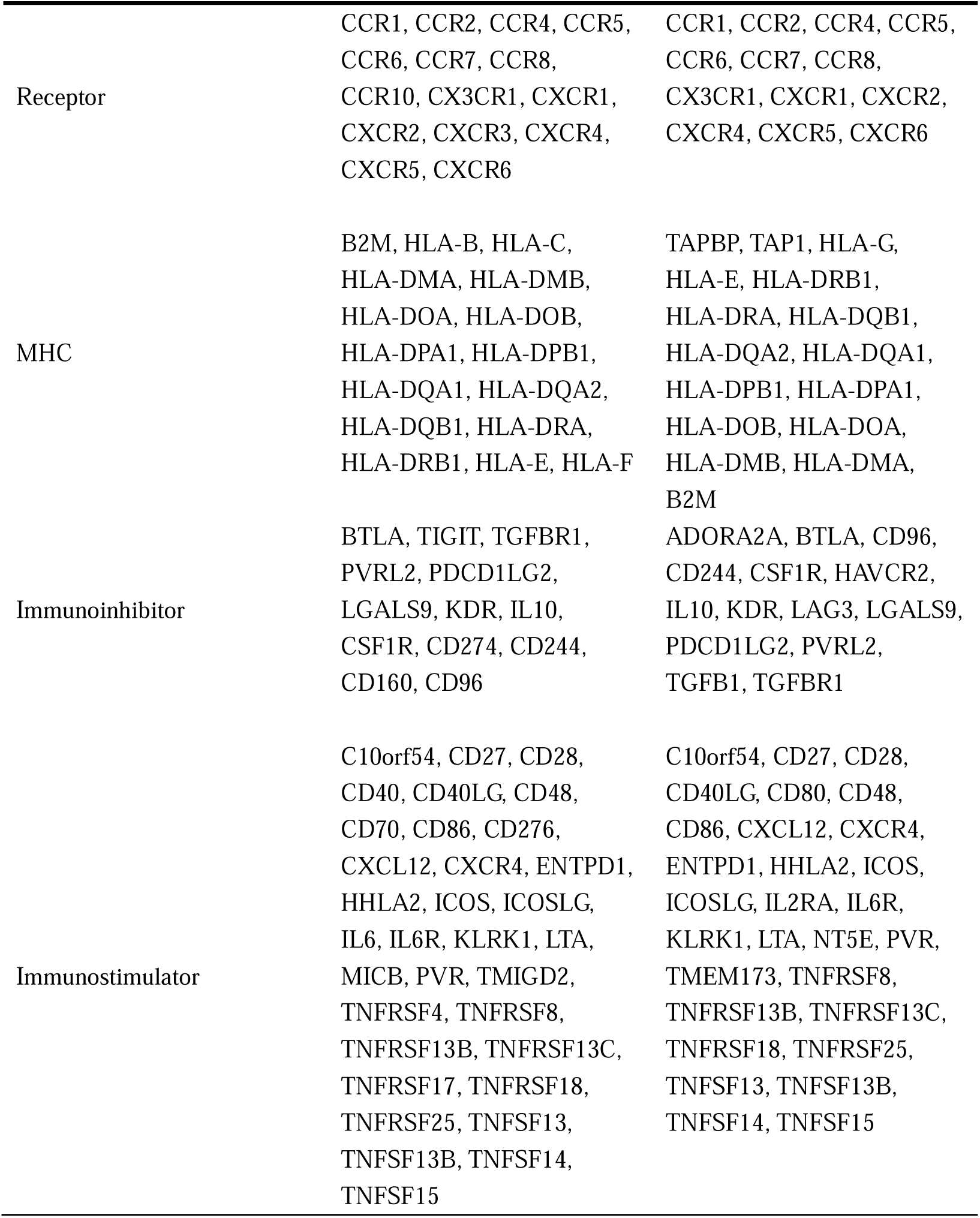
Immunomodulatory genes associated with AChE and BChE Correlation ACHE BCHE.

| Correlation | ACHE | BChE |
| --- | --- | --- |
| Chemokine | XCL2, XCL1, CXCL16,<br>CXCL13, CXCL12, CXCL8,<br>CXCL3, CXCL2, CXCL1,<br>CCL26, CCL24, CCL23,<br>CCL22, CCL20, CCL19,<br>CCL17, CCL15, CCL14,<br>CCL7, CCL5 | CCL2, CCL7, CCL13,<br>CCL14, CCL15, CCL17,<br>CCL18, CCL19, CCL20,<br>CCL21, CCL22, CCL23,<br>CCL24, CCL26, CX3CL1,<br>CXCL6, CXCL12, CXCL14,<br>CXCL16 |
| Receptor | CCR1, CCR2, CCR4, CCR5, CCR6, CCR7, CCR8, CCR10, CX3CR1, CXCR1, CXCR2, CXCR3, CXCR4, CXCR5, CXCR6 | CCR1, CCR2, CCR4, CCR5, CCR6, CCR7, CCR8, CX3CR1, CXCR1, CXCR2, CXCR4, CXCR5, CXCR6 |
| MHC | B2M, HLA-B, HLA-C, HLA-DMA, HLA-DMB, HLA-DOA, HLA-DOB, HLA-DPA1, HLA-DPB1, HLA-DQA1, HLA-DQA2, HLA-DQB1, HLA-DRA, HLA-DRB1, HLA-E, HLA-F | TAPBP, TAP1, HLA-G, HLA-E, HLA-DRB1, HLA-DRA, HLA-DQB1, HLA-DQA2, HLA-DQA1, HLA-DPB1, HLA-DPA1, HLA-DOB, HLA-DOA, HLA-DMB, HLA-DMA, B2M |
| Immunoinhibitor | BTLA, TIGIT, TGFBR1, PVRL2, PDCD1LG2, LGALS9, KDR, IL10, CSF1R, CD274, CD244, CD160, CD96 | ADORA2A, BTLA, CD96, CD244, CSF1R, HAVCR2, IL10, KDR, LAG3, LGALS9, PDCD1LG2, PVRL2, TGFB1, TGFBR1 |
| Immunostimulator | C10orf54, CD27, CD28, CD40, CD40LG, CD48, CD70, CD86, CD276, CXCL12, CXCR4, ENTPD1, HHLA2, ICOS, ICOSLG, IL6, IL6R, KLRK1, LTA, MICB, PVR, TMIGD2, TNFRSF4, TNFRSF8, TNFRSF13B, TNFRSF13C, TNFRSF17, TNFRSF18, TNFRSF25, TNFSF13, TNFSF13B, TNFSF14, TNFSF15 | C10orf54, CD27, CD28, CD40LG, CD80, CD48, CD86, CXCL12, CXCR4, ENTPD1, HHLA2, ICOS, ICOSLG, IL2RA, IL6R, KLRK1, LTA, NT5E, PVR, TMEM173, TNFRSF8, TNFRSF13B, TNFRSF13C, TNFRSF18, TNFRSF25, TNFSF13, TNFSF13B, TNFSF14, TNFSF15 |

**Figure 4.**
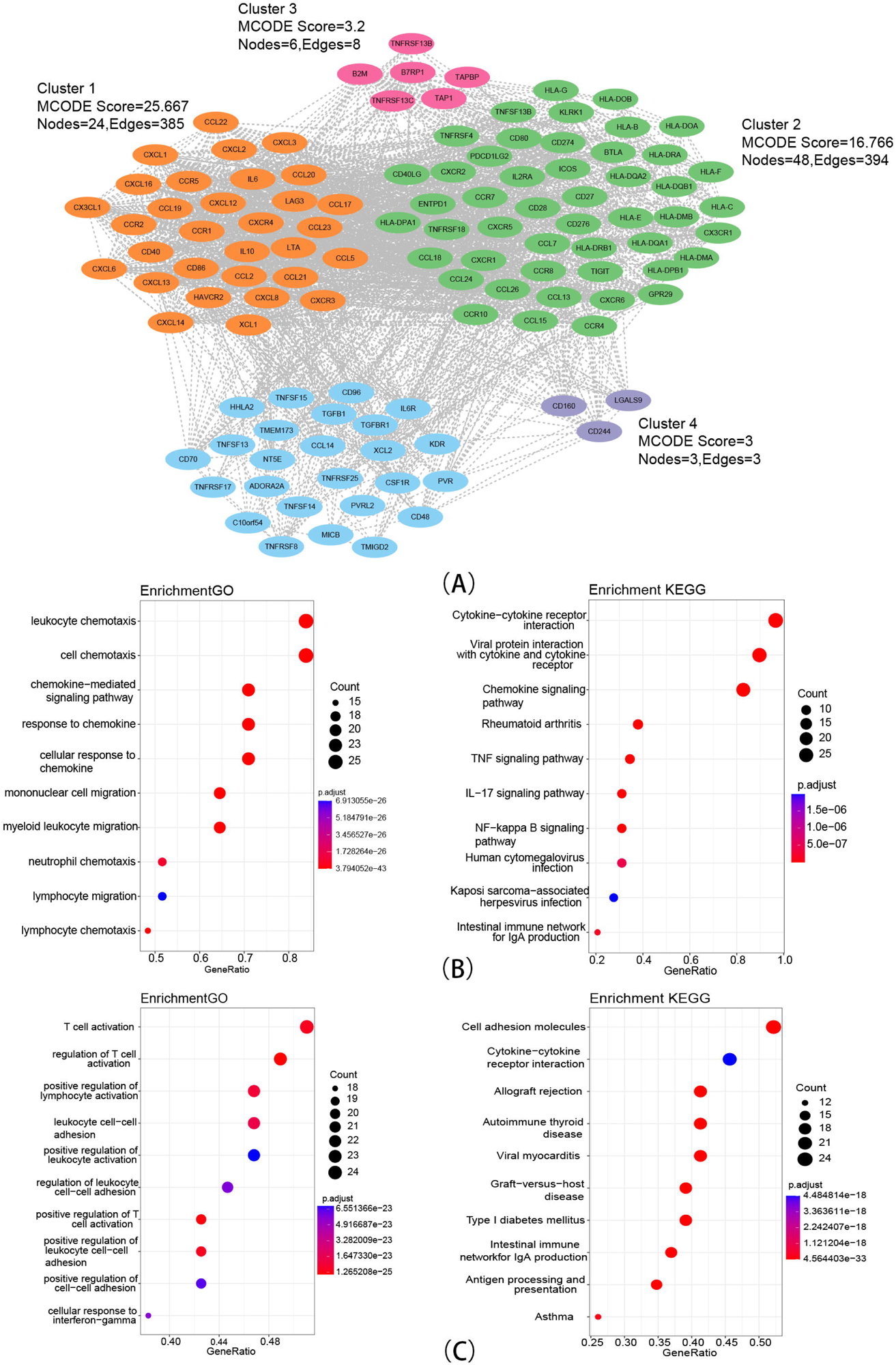
The immunomodulatory networks related to AChE and BChE. (A) PPI network composing of 113 immune genes and four clusters; (B) functional enrichment results of genes in cluster 1; (C) functional enrichment results of genes in cluster 2.

### 3.5 TIDE scores analysis of AChE and BChE

We found that TIDE scores in the AChE high expression group were markedly lower than those in the AChE low expression group, suggesting that LUAD patients with high AChE expression in LUAD may have better outcomes with immunotherapy (Figure 5). AChE expression was negatively correlated with Dysfunction score, while it was not significantly correlated with Exclusion score, indicating that some CD8+T cells in the low expression group were dysfunctional. Since there was no significant difference in CD8+T cell content between the high and low AChE expression group (Figure 3), the worse immune outcome of patients in the low AChE expression group could be related to CD8+T cells dysfunction, which was favorable for cancer cells to undergo immune escape. Similarly, some CD8+T cells in the BChE high expression group were dysfunctional, leading to a possible better outcome of LUAD patients in the BChE low expression group receiving immunotherapy.

**Figure 5.**
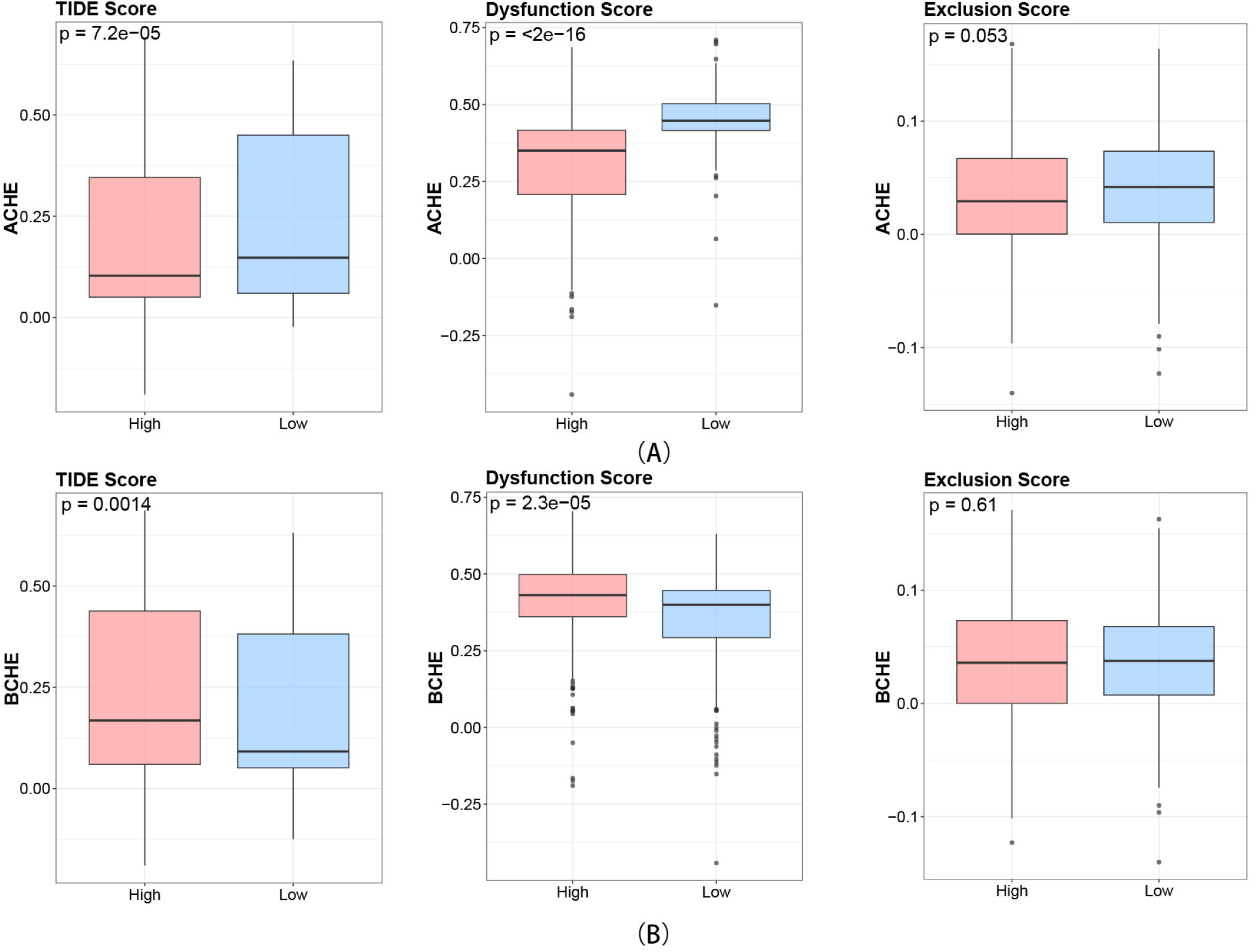
The immunotherapy response related to AChE and BChE. The relationship between AChE, BChE expression and TIDE score, Dysfunction score and Exclusion score.

## 4. Discussion

The cholinergic system exists in both neuronal and non-neuronal cells and forms a complex network that plays essential roles in various physiological functions (Halder and Lal 2021). The main components of the cholinergic system includes the neurotransmitter ACh, AChRs, mAChRs, choline acetyltransferase (CHAT), AChE and BChE, etc..In neurons, ACh is synthesized from choline and acetyl-CoA under the catalysis of CHAT, and is subsequently transported into vesicles and released into the synaptic gap, where it binds to nAChRs or mAChRs to transmit the action potential. Within the synapse, excess ACh is rapidly broken down into choline and acetate by AChE or BChE, thus terminating the excitatory effect of ACh on the postsynaptic membrane. Then, choline is transported back to the cytoplasm and a new cycle of acetylcholine synthesis and release begins (Friedman et al. 2019). Besides neurons, other cells are also able to synthesize ACh (Rosas-Ballina et al. 2011; Takahashi et al. 2014; Wessler and Kirkpatrick 2008).

The cholinergic system plays an important role in the growth and progression of different types of cancers, and has become promising target for cancer therapy (Arese et al. 2018; Halder and Lal 2021; Hutchings et al. 2020; Paleari et al. 2008; Song et al. 2008; Spindel 2016). Studies have shown that many components of the cholinergic system are abnormally expressed in LUAD and other types of lung cancer, To explore the potential role of the cholinergic system in LUAD, we analyzed the mRNA expression profiles of cholinergic genes in LUAD samples and screened 37 genes differentially expressed between the tumor tissues and normal tissues. Among these genes, seven were related to the prognosis of LUAD, including two acetylcholine hydrolase, AChE and BChE.

We found that the elevated expression levels of AChE and BChE were associated with good prognosis of LUAD. In addition to its role in synaptic transmission, AChE can regulate cell proliferation, apoptosis and metastasis (Jiang et al. 2020; Richbart et al. 2021). Decreased cholinesterases activity may enhance cholinergic signaling thereby promoting lung cancer progression (Martinez-Moreno et al. 2006). As an attractive biomarker in cancer diagnosis, BChE has been demonstrated to regulate the development of prostate and pancreatic cancers (Gu et al. 2018; Klocker et al. 2020). Decreased plasma BChE has been associated with poor prognosis in multiple cancer types (Santarpia et al. 2013). By analyzing clinic-pathological factors, we found that the high AChE expression was correlated with good prognosis in male LUAD patients. BChE expression is positively correlated with good prognosis in female patients and non-smoking patients, and the prognosis was best in female non-smoking patients. However, the precise mode of action of BChE in female non-smoking patients is still unclear. The association between BChE and prognosis in female non-smoking patients may facilitate a better understanding of the pathogenesis of non-smoking LUAD patients and provide new ideas for the prevention, diagnosis and treatment of LUAD.

Cancer is considered to be an evolving dynamic process, and the interaction between the tumor microenvironment (TME) and tumor cells plays a key role in tumor development, invasion and metastasis, and treatment (Xiao and Yu 2021). The tumor microenvironment includes immune cells, stromal cells, blood vessels, and extracellular matrix. The tumor microenvironment has attracted much attention as a promising target for therapeutic intervention in tumors (Anderson and Simon 2020). The cholinergic system modulates components of the tumor microenvironment to influence the recruitment of immune cells in the TME and the formation of tumor blood vessels. However, specific insights into how cholinesterase regulates tumor microenvironment components and immune pathways related to LUAD are currently lacking. Understanding the relationship between AChE, BChE and immune microenvironment can help us better understand the effects of AChE and BChE on LUAD (Bader et al. 2020). Results indicated a higher proportion of immune cells in the AChE and BChE high-expression groups compared to the low-expression group, suggesting that the tumor growth and metastasis process was more likely to be influenced by immune cells in the high-expression group. In addition, tumor-associated immune cells play a major role in regulating immune responses. The number of stromal cells was also significantly higher in the BChE high expression group than in the low expression group, and stromal cells are closely associated with drug resistance and immune escape, which also provides new clues for the study of BChE-related cancer drugs (Park et al. 2020). Besides, tumor purity is a potential prognostic tumor marker (Yadav and De 2015), and we found that both AChE and BChE expressions were negatively correlated with tumor purity in LUAD.

We found that monocytes, M0 microphages were associated with good prognosis of lung adenocarcinoma, and M2 microphages, dendritic cells, gamma delta T cells, and regulatory T cells were associated with poor prognosis. Based on the results from TIMER and CIBERSORT, we found that AChE and BChE could regulate immune cells to inhibit the development of lung adenocarcinoma. The abundance of M0 microphages was significantly higher in the AChE high expression group than in the low expression group, and M2 microphages and dendritic cells were more abundant in the low expression group; the abundance of monocytes and M0 microphages was considerably higher in the BChE high expression group than in the low expression group, and regulatory T cells was more abundant in the low expression group. Monocytes are important regulators in the development of cancer, and they play the dual role of either promoting cell survival or promoting cell death, depending on the subpopulation. In the prognostic analysis, monocytes were related to a high survival rate, which may be related to the fact that monocytes can recruit NK cells to kill tumor cells (Li et al. 2018). Moreover, monocytes can differentiate into macrophages and dendritic cells, which may also contribute to the higher M0 microphages content in the BChE high expression group (Goudot et al. 2017). The results of Chen et al. (Chen et al. 2021) are consistent with our findings that M2 microphages can promote lung adenocarcinoma cell invasion, migration and tumor metastasis in vivo. Dendritic cells can present antigens and trigger T-cell mediated immune responses. Interestingly, our results revealed that dendritic cells are associated with poor prognosis. We speculate that dendritic cells are defective in differentiation and activation, resulting in dendritic cell dysfunction that cannot activate anti-tumor immunity, thereby favoring tumor development (Veglia and Gabrilovich 2017). Regulatory T cells suppress the activation of the immune system and contribute to the immune escape of lung cancer, which confirms our results (Zhang et al. 2015). Notably, AChE expression was negatively correlated with CD8+ T content in TIMER results, but the difference in CD8+ T cell content between the two groups with high and low AChE expression was not significant, and the reason for this difference may be related to the number of samples, tumor stage and other factors.CD8+ T cells play a central role in mediating anti-tumor immunity. In order to understand the impact of AChE on tumor development, further research on AChE and CD8+ T cells is necessary.

Immunomodulatory genes closely related to AChE and BChE mainly act on chemokines, cytokines, and T-cell activation pathways, suggesting that AChE and BChE may influence the development of lung adenocarcinoma by regulating these pathways. Chemokines have been proven to be tightly related to the immune landscape of TME and may be good candidates for predicting immunotherapeutic response(Ozga et al. 2021). We also analyzed the relationship between AChE, BChE and immunotherapy response, and the results suggest that patients in the AChE high-expression group and BChE low-expression group may benefit from immunotherapy.Meanwhile, the immune dysregulation of CD8+T cells may be the reason why the AChE low expression group and the BChE high expression group are more prone to immune escape. Taken together, these data provide new insights into a better understanding of the mechanism of action between the cholinergic network and the immune system, as well as cancer therapy.

Although the results of this preliminary study are quite encouraging, it still has some limitations. Firstly, we mainly analyzed only cholinesterases and did not analyze other important proteins in the cholinergic pathway, which has not allowed us to fully understand the mechanism of action of the cholinergic pathway on LUAD. In addition, we only performed bioinformatic analysis and did not validate the tumor suppressive effects of AChE and BChE in LUAD using molecular biology methods. Therefore, further research is needed for a more extensive and comprehensive analysis of the effects of cholinergic pathways on tumor development.

## 5. Conclusion

The cholinergic system can affect the development of LUAD, and our results suggest that cholinesterases AChE and BChE are able to exert an inhibitory effect on LUAD by affecting the tumor microenvironment and modulating immune pathways. Thus, they may be promising therapeutic target for LUAD treatment.

## Data availability statement

The original contributors presented in the study are included in the article/Supplementary Material, further inquiries can be directed to the corresponding author.

## Acknowledgments

The authors would like to thank Dr. Pan Guo for the help with the data collection in study.

