## Supplementary figures and images for "Analyzing the Involvement of Cholinesterases in the Immune Landscape of Lung Adenocarcinoma and Their Prognostic Values"

### Supplemental Figure S1

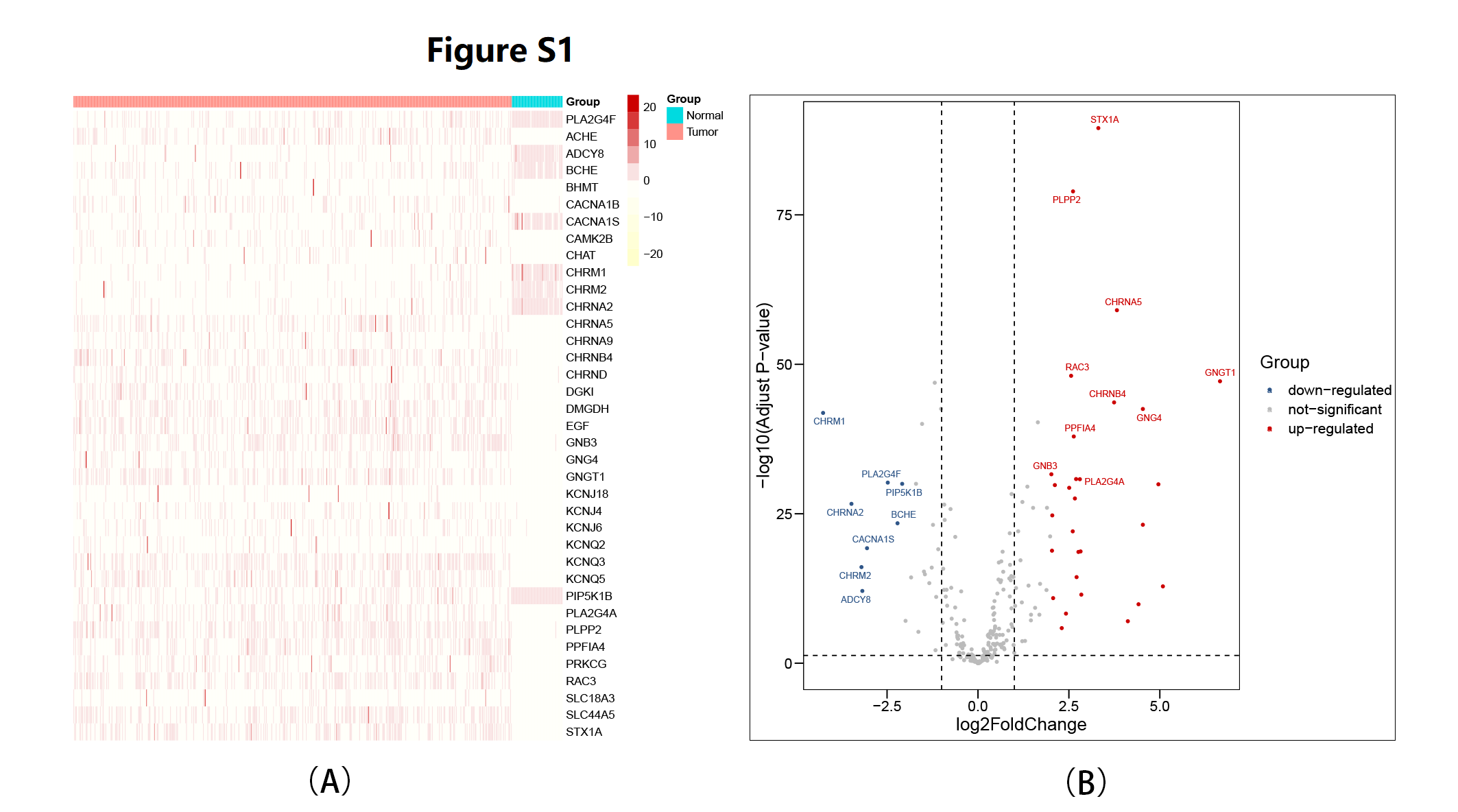

### Supplemental Figure S2

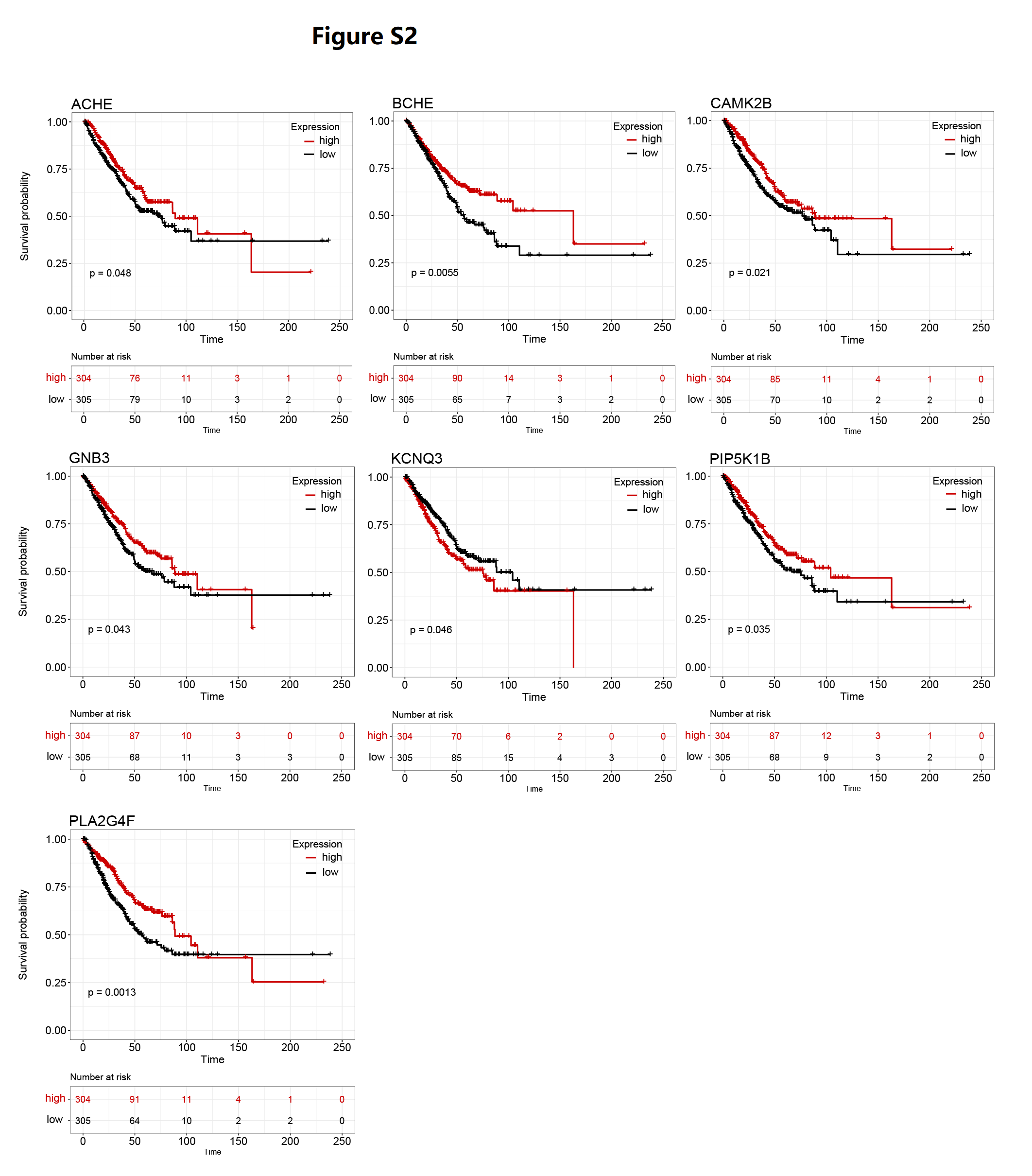

### Supplemental Figure S3

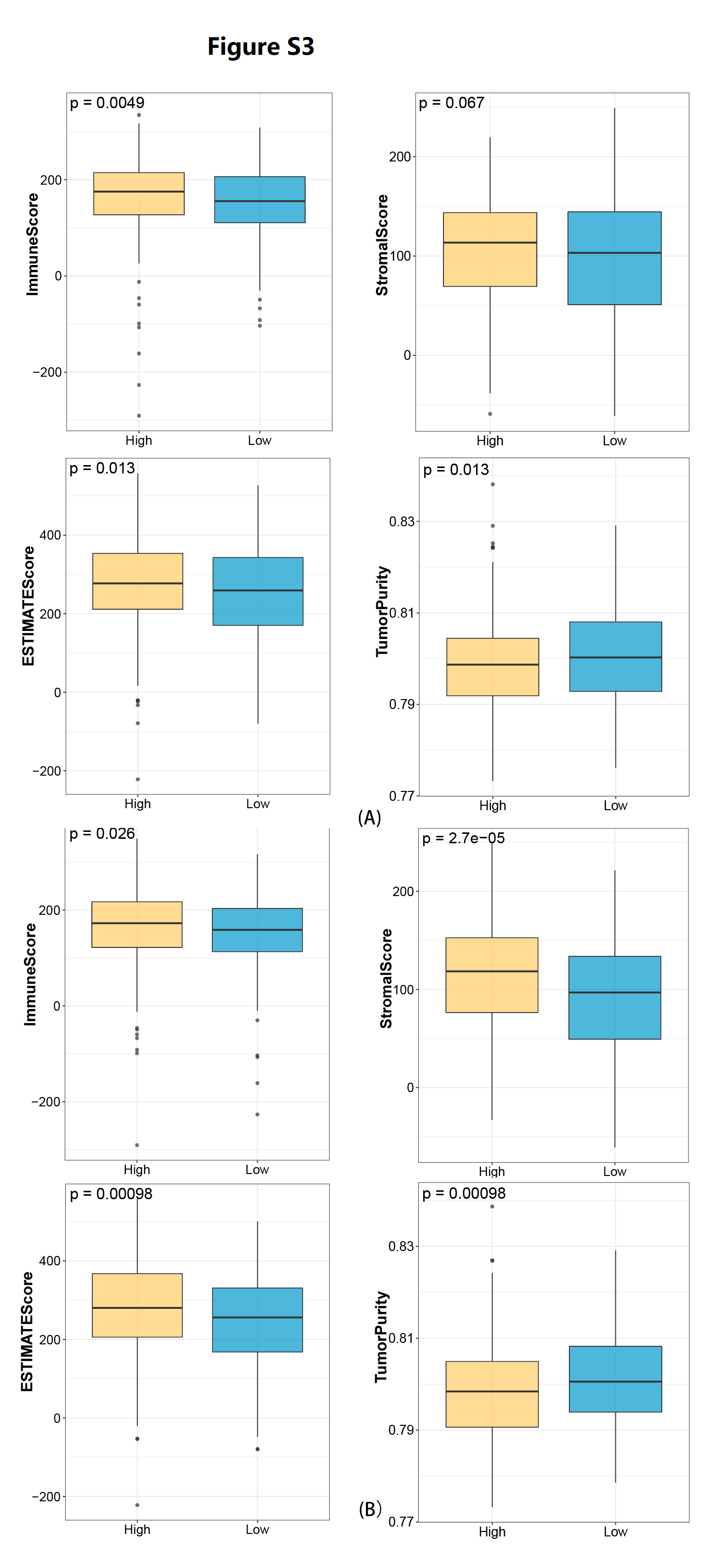

### Supplemental Figure S4

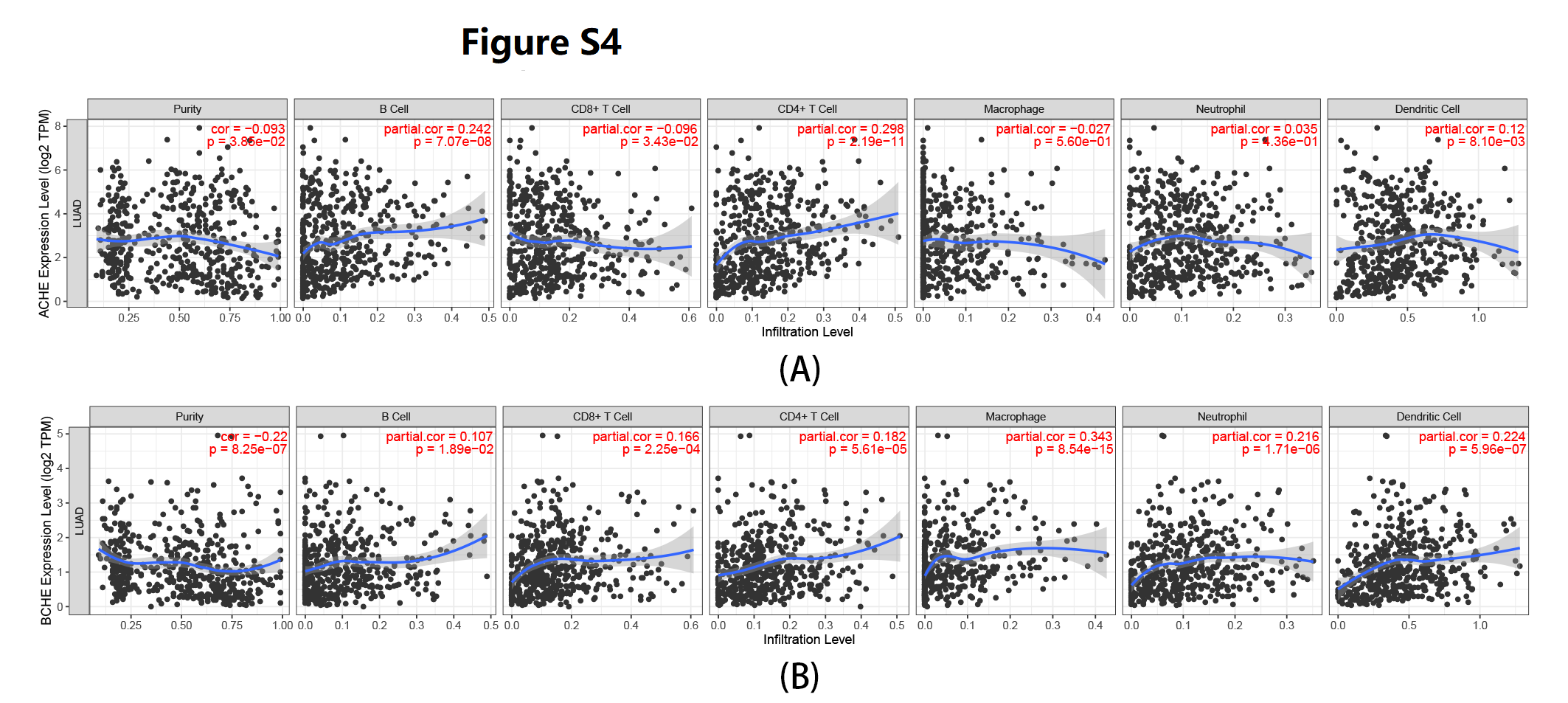

### Supplemental Figure S5

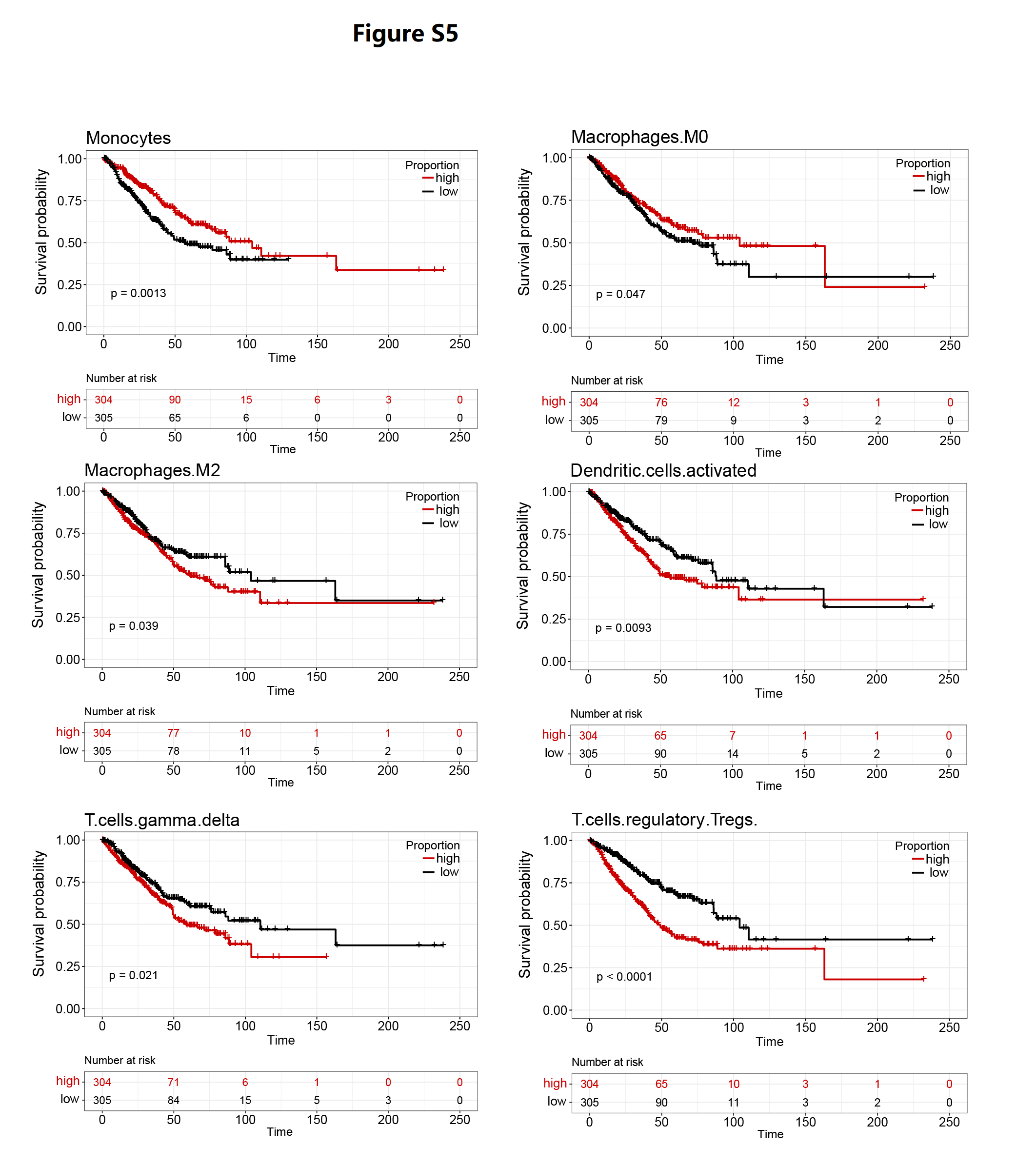
