## Supplemental Tables for "Analyzing the Involvement of Cholinesterases in the Immune Landscape of Lung Adenocarcinoma and Their Prognostic Values"

Table S1. Clinical-pathological information of LUAD patients

|  | TCGA^a^ | GEO^b^ | Total^c^ |
| --- | --- | --- | --- |
| Gender | Male:Female = 227:267 | Male:Female=17:98 | Male:Female=244:365 |
| Age | 30~50：38  51~70：287  71~90：159  NA:10 | 30~50：9  51~70：104  71~90：2 | 30~50：47  51~70：391  71~90：161  NA:10 |
| Stage | I:II:III:IV=267:120:79:26  NA=2 | I:II=94:21 | I:II:III:IV = 361:141:79:26  NA=2 |
| Smoking status | Smoker:Nonsmoker = 380:114 | Nonsmoker=115 | Smoker:Nonsmoker = 380:229 |
| Race | Asian:White:Other = 7:381:53  NA=53 | NA | Asian:White:Other = 7:381:53  NA=168 |
| Survival Status | Alive:Dead = 315:179 | Alive:Dead = 100:15 | Alive:Dead = 415:194 |
| Patients | 494 | 115 | 609 |

^a^ LUAD samples retrieved from the TCGA database.

^b^ LUAD samples retrieved from the dataset GSE31210 in GEO database.

^c^ LUAD samples from TCGA and GEO were merged.

Table S2. The impact of clinical features on prognosis in LUAD patients

| Characteristics | Univariate Cox | | | Multivariate Cox | | |
| --- | --- | --- | --- | --- | --- | --- |
|  | Hazard Ratio | 95%CI | P-value | Hazard Ratio | 95%CI | P-value |
| Gender | 0.73 | 0.55-0.97 | 0.028 | 0.92 | 0.69-1.2 | 0.59 |
| Age | 1.02 | 1.01-1.04 | 0.004 | 1 | 0.99-1 | 0.0056 |
| Stage | 1.98 | 1.73-2.27 | 0.001 | 1.8 | 1.6-2.1 | 3.90E-16 |
| Smoking status | 0.44 | 0.32-0.6 | 0.001 | 0.54 | 0.39-0.76 | 0.00029 |
| Race | 0.85 | 0.66-1.09 | 0.196 |  |  |  |

Table S3. Cholinergic pathway-related genes

| Pathway | Gene |
| --- | --- |
| Cholinergic Synapse  （from：KEGG Database） | CHAT, ACHE, SLC18A3, CHRM1, CHRM3, CHRM5, GNAQ, GNA11, PLCB1, PLCB2, PLCB3, PLCB4, ITPR1, ITPR2, ITPR3, PRKCA, PRKCB, PRKCG, KCNQ1, KCNQ2, KCNQ3, KCNQ4, KCNQ5, KCNJ2, KCNJ12, KCNJ18, KCNJ4, KCNJ14, CHRM2, CHRM4, GNAI1, GNAI3, GNAI2, GNA01, GNB1, GNB2, GNB3, GNB4, GNB5, GNG2, GNG3, GNG4, GNG5, GNG7, GNG8, GNG10, GNG11, GNG12, GNG13, GNGT1, GNGT2, KCNJ3, KCNJ6, PIK3CG, PIK3R5, PIK3R6, HRAS, KRAS, NRAS, MAP2K1, MAPK1, MAPK3, FOS, CHRNA7, CHRNA4, CHRNB2, CHRNA3, CHRNB4, CHRNA6, ADCY1, ADCY2, ADCY3, ADCY4, ADCY5, ADCY6, ADCY7, ADCY8, ADCY9, PRKACA, PRKACB, PRKACG, CREB1, ATF4, CREB3, CREB3L1, CREB3L2, CREB3L3, CREB3L4, CREB5, CAMK2A, CAMK2D, CAMK2B, CAMK2G, CAMK4, JAK2, FYN, PIK3CA, PIK3CB, PIK3CD, PIK3R1, PIK3R2, PIK3R3, AKT1, AKT2, AKT3, BCL2, CACNA1A, CACNA1B, CACNA1C, CACNA1D, CACNA1F, CACNA1S, SLC5A7 |
| Choline Metabolism in Cancer  （from：KEGG Database） | EGF, PDGFA, PDGFB, PDGFC, PDGFD, EGFR, PDGFRA, PDGFRB, GRB2, SOS1, SOS2, HRAS, KRAS, NRAS, RAF1, MAP2K1, MAP2K2, MAPK1, MAPK3, RALGDS, MAPK8, MAPK10, MAPK9, PLA2G4E, PLA2G4A, JMJD7-PLA2G4B, PLA2G4B, PLA2G4C, PLA2G4D, PLA2G4F, PIK3CA, PIK3CD, PIK3CB, PIK3R1, PIK3R2, PIK3R3, PDPK1, AKT1, AKT2, AKT3, TSC1, TSC2, RHEB, MTOR, RPS6KB1, RPS6KB2, EIF4EBP1, PIP5K1C, PIP5K1A, PIP5K1B, WAS, RAC1, RAC2, RAC3, WASF1, WASF2, WASF3, SP1, PLD1, PLD2, SLC5A7, SLC44A1, SLC44A4, SLC44A5, SLC44A2, SLC44A3, SLC22A1, SLC22A2, SLC22A3, SLC22A5, SLC22A4, CHKA, CHKB, HIF1A, JUN, FOS, PCYT1B, PCYT1A, CHPT1, PLCG1, PLPP1, PLPP3, PLPP2, DGKZ, DGKD, DGKI, DGKA, DGKE, DGKB, DGKH, DGKG, DGKQ, DGKK, PRKCA, PRKCB, PRKCG, LYPLA1, GPCPD1 |
| Choline catabolism  （from：Reactome Pathway Database）  Acetylcholine Neurotransmitter Release Cycle  （from：Reactome Pathway Database） | DMGDH, SLC44A1, ALDH7A1, BHMT, SARDH, CHDH  STXBP1, CPLX1, RIMS1, SNAP25, RAB3A, SYT1, VAMP2, STX1A, TSPOAP1, UNC13B, PPFIA2, PPFIA4, PPFIA3, PPFIA1, SLC18A3, CHAT, SLC5A7 |
| Choline Biosynthesis III  （from：Pathway Commons Database） | PCYT1B, PLD3, PCYT1A, CEPT1, CHPT1, PLD4, PLD6, PLD2, PLD1 |
| Acetylcholine Synthesis  （from：WikiPathways Database） | ACHE, CHAT, CHKA, PCYT1A, PDHA1, PDHA2, PEMT |
| Others  （from：PubMed） | SLC18A3, SNCA, SLC5A7, CHAT, SLC18A3, SLC44A1, SLC10A4, SLC5A7, SLC44A1, SLC44A2, SLC44A3, SLC44A4, SLC44A5, CHRM1, CHRM2, CHRM3, CHRM4, CHRM5, CHRNA1, CHRNA2, CHRNA3, CHRNA4, CHRNA5, CHRNA6, CHRNA7, CHRNA9, CHRNA10, CHRNB1, CHRNB2, CHRNB3, CHRNB4, CHRNG, CHRND, CHRNE, ACHE, BCHE, CRAT |

Table S4. 37 differentially expressed genes

| Symbol | Gene name | logFC | Pval | adj.Pval | Group |
| --- | --- | --- | --- | --- | --- |
| STX1A | syntaxin 1A | 3.309519 | 1.33E-92 | 3.06E-90 | up-regulated |
| PLPP2 | phospholipid phosphatase 2 | 2.613459 | 1.03E-81 | 1.19E-79 | up-regulated |
| CHRNA5 | cholinergic receptor nicotinic alpha 5 subunit | 3.820368 | 1.16E-61 | 8.93E-60 | up-regulated |
| RAC3 | Rac family small GTPase 3 | 2.560681 | 1.45E-50 | 8.35E-49 | up-regulated |
| GNGT1 | G protein subunit gamma transducin 1 | 6.654972 | 1.48E-49 | 6.85E-48 | up-regulated |
| CHRNB4 | cholinergic receptor nicotinic beta 4 subunit | 3.74297 | 7.14E-46 | 2.36E-44 | up-regulated |
| GNG4 | G protein subunit gamma 4 | 4.533805 | 1.16E-44 | 2.98E-43 | up-regulated |
| CHRM1 | cholinergic receptor muscarinic 1 | -4.25921 | 5.85E-44 | 1.35E-42 | Down-regulated |
| PPFIA4 | PTPRF interacting protein alpha 4 | 2.63381 | 6.68E-40 | 1.19E-38 | up-regulated |
| GNB3 | G protein subunit beta 3 | 2.017645 | 1.50E-33 | 2.48E-32 | up-regulated |
| PLA2G4A | phospholipase A2 group IVA | 2.697886 | 1.02E-32 | 1.57E-31 | up-regulated |
| KCNQ5 | potassium voltage-gated channel subfamily Q member 5 | 2.801883 | 1.15E-32 | 1.66E-31 | up-regulated |
| PLA2G4F | phospholipase A2 group IVF | -2.48063 | 4.54E-32 | 6.17E-31 | down-regulated |
| PIP5K1B | phosphatidylinositol-4-phosphate 5-kinase type 1 beta | -2.08443 | 8.17E-32 | 9.93E-31 | down-regulated |
| CHRNA9 | cholinergic receptor nicotinic alpha 9 subunit | 4.960961 | 1.03E-31 | 1.18E-30 | up-regulated |
| KCNQ3 | potassium voltage-gated channel subfamily Q member 3 | 2.113985 | 1.46E-31 | 1.60E-30 | up-regulated |
| EGF | epidermal growth factor | 2.507978 | 4.56E-31 | 4.58E-30 | up-regulated |
| ACHE | acetylcholinesterase | 2.66494 | 3.00E-29 | 2.77E-28 | up-regulated |
| CHRNA2 | cholinergic receptor nicotinic alpha 2 subunit | -3.47903 | 2.57E-28 | 2.20E-27 | down-regulated |
| DGKI | diacylglycerol kinase iota | 2.043127 | 2.61E-26 | 1.89E-25 | up-regulated |
| BCHE | butyrylcholinesterase | -2.21285 | 5.75E-25 | 3.91E-24 | down-regulated |
| KCNQ2 | potassium voltage-gated channel subfamily Q member 2 | 4.53382 | 1.05E-24 | 6.92E-24 | up-regulated |
| SLC44A5 | solute carrier family 44 member 5 | 2.605519 | 1.44E-23 | 8.74E-23 | up-regulated |
| CACNA1S | calcium voltage-gated channel subunit alpha1 S | -3.05108 | 1.05E-20 | 5.77E-20 | down-regulated |
| DMGDH | dimethylglycine dehydrogenase | 2.035078 | 2.87E-20 | 1.51E-19 | up-regulated |
| PRKCG | protein kinase C gamma | 2.820787 | 3.92E-20 | 2.01E-19 | up-regulated |
| CAMK2B | calcium/calmodulin dependent protein kinase II beta | 2.761975 | 4.97E-20 | 2.44E-19 | up-regulated |
| CHRM2 | cholinergic receptor muscarinic 2 | -3.20389 | 1.89E-17 | 8.38E-17 | down-regulated |
| BHMT | betaine--homocysteine S-methyltransferase | 2.711335 | 1.03E-15 | 3.98E-15 | up-regulated |
| SLC18A3 | solute carrier family 18 member A3 | 5.084896 | 4.23E-14 | 1.40E-13 | up-regulated |
| ADCY8 | adenylate cyclase 8 | -3.18016 | 2.73E-13 | 8.20E-13 | down-regulated |
| CACNA1B | calcium voltage-gated channel subunit alpha1 B | 2.84108 | 1.17E-12 | 3.38E-12 | up-regulated |
| KCNJ6 | potassium inwardly rectifying channel subfamily J member 6 | 2.06686 | 4.73E-12 | 1.30E-11 | up-regulated |
| CHAT | choline O-acetyltransferase | 4.414656 | 5.20E-11 | 1.38E-10 | up-regulated |
| CHRND | cholinergic receptor nicotinic delta subunit | 2.419562 | 2.09E-09 | 5.03E-09 | up-regulated |
| KCNJ18 | potassium inwardly rectifying channel subfamily J member 18 | 4.120841 | 4.31E-08 | 9.49E-08 | up-regulated |
| KCNJ4 | potassium inwardly rectifying channel subfamily J member 4 | 2.303366 | 6.51E-07 | 1.35E-06 | up-regulated |
